# Cyclin B3 activates the Anaphase-Promoting Complex/Cyclosome in meiosis and mitosis

**DOI:** 10.1101/2020.06.05.136291

**Authors:** Damien Garrido, Mohammed Bourouh, Éric Bonneil, Pierre Thibault, Andrew Swan, Vincent Archambault

## Abstract

In mitosis and meiosis, chromosome segregation is triggered by the Anaphase-Promoting Complex/Cyclosome (APC/C), a multi-subunit ubiquitin ligase that targets proteins for degradation, leading to the separation of chromatids. APC/C activation requires phosphorylation of its APC3 and APC1 subunits, which allows the APC/C to bind its Cdc20 co-activator. The identity of the kinase(s) responsible for APC/C activation in vivo is unclear. Cyclin B3 is required for meiotic anaphase in flies, worms and vertebrates, but whether it activates the APC/C is unclear. We found that *Drosophila* Cyclin B3 (CycB3) collaborates with PP2A-B55/Tws in embryonic development, indicating that CycB3 also promotes anaphase in mitosis. Moreover, CycB3 promotes APC/C activity and anaphase in cells in culture. We show that CycB3 physically associates with the APC/C, is required for phosphorylation of APC3, and promotes APC/C association with its co-activators. We propose that CycB3-Cdk1 directly phosphorylates the APC/C to activate it in both meiosis and mitosis.

## INTRODUCTION

Mitosis and meiosis (collectively referred to as M-phase) are distinct modes of nuclear division resulting in diploid or haploid products, respectively. In animals, both require the breakdown of the nuclear envelope, the condensation of chromosomes and their correct attachment on a microtubule-based spindle, where chromosomes are under tension and chromatids are held together by cohesins. Progression through these initial phases requires multiple phosphorylation events of various protein substrates by mitotic kinases including Cyclin-Dependent Kinases (CDKs) activated by their mitotic cyclin partners (Morgan, 2007). M-phase completion from this point (mitotic exit) requires the degradation of mitotic cyclins, and the dephosphorylation of several mitotic phosphoproteins by phosphatases including Protein Phosphatase 2A (PP2A) (Holder et al., 2019). Mitotic exit begins with the segregation of chromosomes in anaphase. In mitosis, sister chromatids segregate. In meiosis I, replicated homologous chromosomes segregate, and in the subsequent meiosis II, sister chromatids segregate. Nuclear divisions are completed with the reassembly of a nuclear envelope concomitant with the decondensation of chromosomes (Schellhaus et al., 2016). How mitosis and meiosis are alike and differ in the molecular mechanisms of their exit programs is not completely understood.

Chromosome segregation is triggered by the Anaphase-Promoting Complex/Cyclosome (APC/C), a multi-subunit E3 ubiquitin ligase (Alfieri et al., 2017; King et al., 1995; Sudakin et al., 1995; Yamano, 2019). By catalysing the addition of ubiquitin chains on the separase inhibitor securin, the APC/C targets it for degradation by the proteasome (Cohen-Fix et al., 1996; Funabiki et al., 1996). As a result, separase cleaves cohesins, allowing separated chromosomes to migrate towards opposing poles of the spindle (Ciosk et al., 1998; Uhlmann et al., 1999). Activation of the APC/C in mitosis requires its recruitment of its co-factor Cdc20 (Fang et al., 1998; Visintin et al., 1997). This recruitment can be prevented by the Spindle-Assembly Checkpoint (SAC), a complex mechanism that allows the sequestration of Cdc20 until all chromosomes are correctly attached on the spindle (Corbett, 2017). Cdc20 binding to the APC/C is also inhibited by its phosphorylation at CDK sites (Alfieri et al., 2017). Phosphatase activity is then required to dephosphorylate Cdc20 and allow its binding of the APC/C for its activation of anaphase (Fujimitsu and Yamano, 2020; Hein et al., 2017; Labit et al., 2012). In addition, phosphorylation of the APC/C itself is required to allow Cdc20 binding (Kramer et al., 2000). Phosphorylation of APC3/Cdc27 and APC1 is key to this process. Phosphorylation of APC3 at CDK sites promotes the subsequent phosphorylation of APC1, inducing a conformational change in APC1 that opens the Cdc20 binding site (Fujimitsu et al., 2016; Zhang et al., 2016). However, the precise identity of the kinase(s) involved in this process *in vivo* is unknown.

At least 3 types of cyclins contribute to M-phase in animals: Cyclins A, B and B3 (Fung and Poon, 2005; Li et al., 2019a). The Cyclin A type (A1 and A2 in mammals) can activate Cdk1 or Cdk2 and is required for mitotic entry, at least in part by allowing the phosphorylation of Cdc20 to prevent its binding and activation of the APC/C (Hein and Nilsson, 2016). This allows mitotic cyclins to accumulate without being ubiquitinated prematurely by the APC/C and degraded. The Cyclin B type (B1 and B2 in mammals) also promotes mitotic entry and is required for mitotic progression by allowing the phosphorylation of several substrates by Cdk1 (Lindqvist et al., 2009). Mammalian Cyclin B3, which can associate with both Cdk1 and Cdk2, is required for meiosis but its contribution to mitosis is less clear in view of its low expression in somatic cells (Gallant and Nigg, 1994; Zhang et al., 2015b). *Drosophila* possesses a single gene for each M-phase cyclin: *CycA* (*Cyclin A*), *CycB* (*Cyclin B*) and *CycB3* (*Cyclin B3*) that collaborate to ensure mitotic progression by activating Cdk1. Genetic and RNAi results suggest that they act sequentially, CycA being required before prometaphase, CycB before metaphase and CycB3 at the metaphase-anaphase transition (McCleland et al., 2009; Yuan and O’Farrell, 2015). CycA is the only essential cyclin, as it is required for mitotic entry (Lehner and O’Farrell, 1989; Reber et al., 2006). *CycB* and *CycB3* mutants are viable, but mutations of *CycB* and *CycB3* are synthetic-lethal, suggesting redundant roles in mitosis (Jacobs et al., 1998). However, mutation of *CycB* renders females sterile due to defects in ovary development, and mutant males are also sterile (Jacobs et al., 1998).

*Drosophila* CycB3 associates with Cdk1 and is required for female meiosis (Jacobs et al., 1998). In *Drosophila*, eggs normally stay arrested in metaphase I of meiosis until egg laying triggers entry into anaphase I and the subsequent meiosis II. However, *CycB3* mutant eggs predominantly stay arrested in meiosis I (Bourouh et al., 2016). In addition, silencing CycB3 expression in early embryos delays anaphase onset during the syncytial mitotic divisions (Yuan and O’Farrell, 2015). Cyclin B3 is also required for anaphase in female meiosis of vertebrates and worms. In mice, RNAi Knock-down of Cyclin B3 in oocytes inhibits the metaphase-anaphase transition in meiosis I (Zhang et al., 2015a). Recently, two groups independently knocked out the Cyclin B3-coding *Ccnb3* gene in mice and found that they were viable but female-sterile due to a highly penetrant arrest in meiotic metaphase I (Karasu et al., 2019; Li et al., 2019b). In *C. elegans*, the closest Cyclin B3 homolog, CYB-3 is required for anaphase in mitosis (Deyter et al., 2010).

How Cyclin B3 promotes anaphase in any system is unknown. One possibility is that it is required for Cdk1 to phosphorylate the APC/C on at least one of its activating subunits, APC3 or APC1. This has not been investigated. Another possibility is that inactivation of Cyclin B3 leads to an early mitotic defect that activates the SAC. This appears to be the case in *C. elegans*, because inactivation of the SAC rescues normal anaphase onset in the absence of CYB-3 (Deyter et al., 2010). However, in *Drosophila*, inactivation of the SAC by the mutation of *mad2* did not eliminate the delay in anaphase onset observed when CycB3 is silenced in syncytial embryos (Yuan and O’Farrell, 2015). Similarly, in mouse oocytes, silencing Mad2 does not rescue the meiotic metaphase arrest upon Cyclin B3 depletion (Zhang et al., 2015a). In other studies, SAC markers on kinetochores did not persist in metaphase-arrested *Ccnb3* KO oocytes, and SAC inactivation by chemical inhibition of Mps1 did not restore anaphase (Karasu et al., 2019; Li et al., 2019b). Finally, it is also possible that Cyclin B3 is required upstream of another event required for APC/C activation, for example the activation of a phosphatase required for Cdc20 dephosphorylation and subsequent recruitment to the APC/C.

In this study, we have investigated how CycB3 promotes anaphase in *Drosophila*. We report several lines of evidence indicating that CycB3 directly activates the APC/C in both meiosis and mitosis.

## RESULTS AND DISCUSSION

### CycB3 functions upstream of the APC/C and collaborates with PP2A-Tws to promote anaphase in meiosis and embryonic mitosis

In addition to the APC/C, another important enzyme for mitotic exit is PP2A-B55 (Holder et al., 2019). We previously conducted a maternal-effect genetic screen for enhancers of mutations in *tws*, which encodes the B55 regulatory subunit of PP2A in *Drosophila*. A genetic interaction between *tws* and *lamin* led us to identify a role of PP2A-Tws/B55 in nuclear envelope reformation after mitosis and meiosis (Mehsen et al., 2018). Embryos from mothers heterozygous for mutations in *tws* and in *lamin*, displayed penetrant nuclear integrity defects. Here, we investigated a genetic interaction between *tws* and *CycB3*, identified in the same screen. We first validated the genetic interaction using different alleles for each gene (Fig 1A). While *CycB3*^*2*^ is a null allele where part of the coding sequence is deleted, *CycB3*^*L6*^ (*CycB3*^*L6540*^) is a hypomorphic allele induced by the insertion of a P-element (Bourouh et al., 2016; Jacobs et al., 1998). *tws*^*aar1*^ and *tws*^*P*^ are hypomorphic alleles caused by P-element insertions (Mayer-Jaekel et al., 1994). We examined the phenotypes of eggs/embryos produced by mothers heterozygous for *CycB3*^*2*^ and *tws*^*aar1*^ (Fig 1B). We stained for microtubules, total DNA and the pericentric DNA of the X-chromosome. Strikingly, most embryos from double-mutant mothers arrest very early in embryonic development, more specifically in metaphase of the first or second mitotic divisions. In addition, a minority of eggs arrest in metaphase of meiosis I, consistent with the known role of CycB3 in promoting anaphase in meiosis. These phenotypes are not observed in embryos from *CycB3*^*2/*+^ or *tws*^*aar1/*+^ mothers, where nuclear divisions proceed normally.

**Figure 1.**
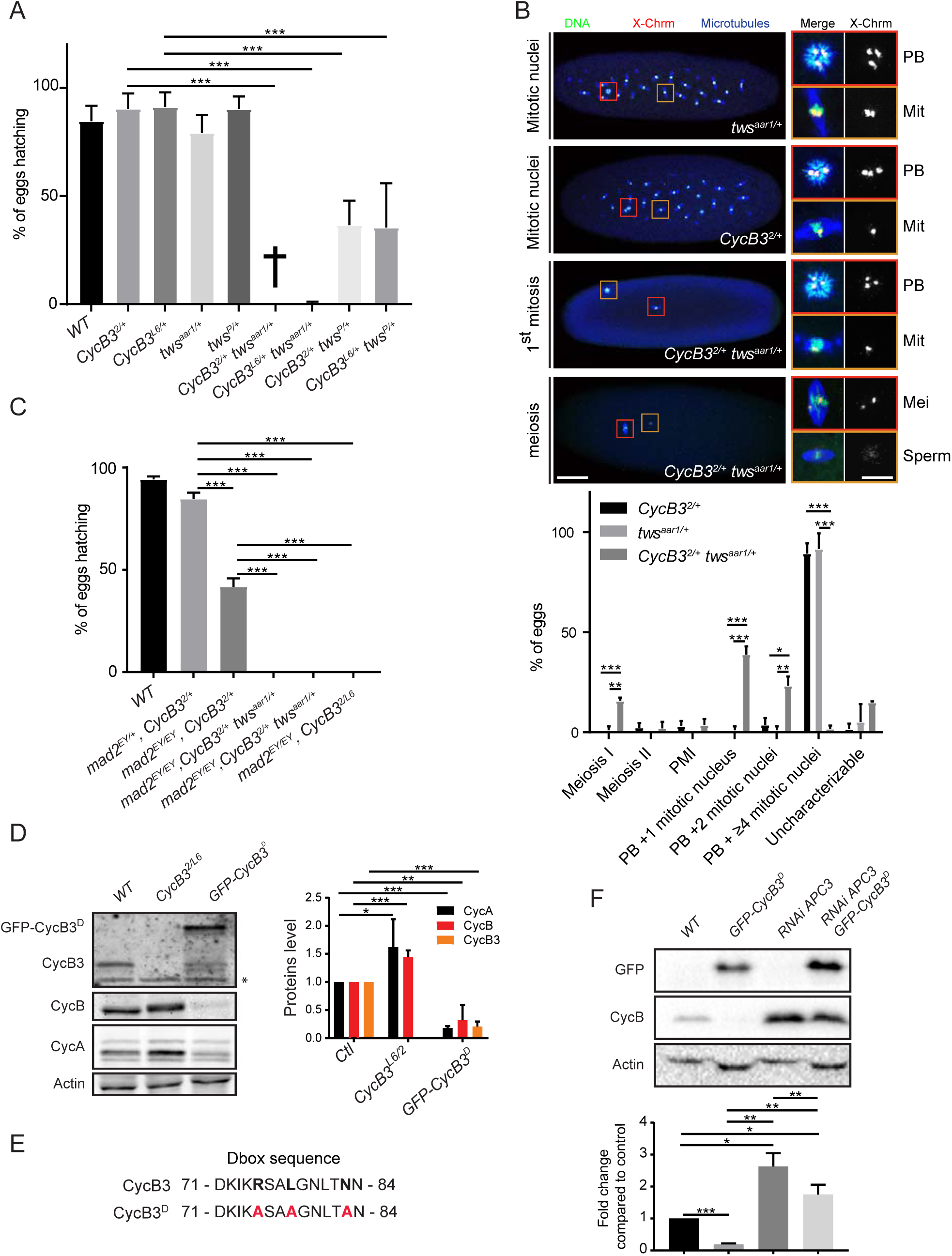
CycB3 collaborates with PP2A-Tws and functions upstream of the APC/C to promote the metaphase-anaphase transition in meiosis and mitosis. A. Mutations in *CycB3* and *tws* enhance each other in a maternal effect. Eggs from mothers of the indicated genotypes were scored for their hatching rate. B. Eggs from *CycB3*^*2/*+^ *tws*^*aar1/*+^ mothers arrest in metaphase of meiosis or mitosis. Green: DNA, Blue: α-Tubulin, Red: FISH for pericentric DNA of the X chromosome. Mitotic spindles (Mit) are recognized by the presence of astral microtubules, co-occurrence with a polar body (PB), and 1 or 2 foci of pericentromeric X-chromosome DNA. Meiotic I metaphase spindles (Mei) are recognized by the absence of astral microtubules, the absence of a polar body in the egg, and at least 2 foci of pericentromeric X-chromosome DNA. In a meiotic arrest, the sperm nucleus can break and nucleate a spindle (bottom images). Insets from the left are enlarged on the right with corresponding color frames. Scale bars: left panel 50 µm, right panel 10 µm. PMI: post-meiotic interphase. C. Inactivation of the SAC by mutation of *mad2* does not rescue the development of embryos from *CycB3*^*2/L6*^ or *CycB3*^*2/*+^ *tws*^*aar1/*+^ mothers. Eggs from mothers of the indicated genotypes were scored for their hatching rate. D. Mutation of *CycB3* results in higher CycA and CycB levels, while maternal expression of GFP-CycB3^D^ results in lower CycA, CycB and endogenous CycB3 levels in eggs. WT (Wild type): non-fertilized eggs. *Non-specific band. E. Mutation of the destruction box motif in CycB3^D^. F. CycB3 negatively regulates CycB levels in an APC-dependent manner. Eggs were collected for 2 hrs from the indicated conditions and analyzed by Western blot. Expression of *UASp-GFP-CycB3*^*D*^ and of *UASp-APC3 RNAi* was driven maternally by *Mat α-Tubulin Gal4-VP16*. The *UASp-GFP*-*CycB3*^*D*^ and *UASp-APC3 RNAi* alone genotypes also contained a *UASp-WHITE* construct, to control for potential dilution of Gal4. Eggs from all conditions failed to develop (WT: unfertilized eggs). Error bars: SD. \*\*\**p* < 0.001; \*\**p* < 0.01; \**p* < 0.05 from ANOVA for panels A and C and from paired *t*-tests for panels B, D and F. See also Figure S1.

To test if the meiotic and mitotic metaphase arrests observed when CycB3 function is compromised are due to activation of the SAC, we introduced a null mutation in *mad2* (*mad2*^*EY21687*^ = *mad2*^*EY*^) known to inactivate the SAC (Li et al., 2010). We found that inactivation of *mad2* does not rescue the development of embryos from *CycB3*^*2/*+^ *tws*^*aar1/*+^ or *CycB3*^*2/L6*^ mothers (Fig 1C). In fact, inactivation of *mad2* tend to enhance hatching defects when CycB3 is partially inactivated. Therefore, the metaphase arrest under a loss of CycB3 function is not caused by a defect that activates the SAC.

Based on results obtained in flies and mice, it has been suggested that CycB3 may function upstream of the APC/C to activate it (Karasu et al., 2019; Li et al., 2019b; Yuan and O’Farrell, 2015; Zhang et al., 2015a). We decided to test this hypothesis. First, we found that levels of CycA and CycB, both targets of the APC/C, are higher in eggs from *CycB3*^*2/L6*^ mothers (Fig 1D). To test if overexpressing a stabilized form of CycB3 would conversely lead to a destabilization of mitotic cyclins, we generated transgenic flies for the expression of CycB3 mutated in its destruction box (Fig 1E) and N-terminally fused to GFP. Expression of GFP-CycB3^D^ in the female germline with Gal4 results in a reduction of CycA, CycB and endogenous CycB3 levels in eggs, which do not hatch (Fig 1D). This result is consistent with an overactivation of the APC/C when CycB3 is stabilized. In this case, the prediction is that inactivation of the APC/C in this context should rescue CycB levels. This is what is observed after RNAi-silencing of APC1 or APC3, both essential subunits of the APC/C (Fig 1F and S1A). Very similar results were obtained with hypomorphic mutations of *cortex* and *fizzy* (Fig S1B), the two Cdc20 co-activators of the APC/C active in female meiosis and syncytial mitoses (Dawson et al., 1995; Pesin and Orr-Weaver, 2007; Swan and Schupbach, 2007). These results confirm that CycB3 functions upstream of the APC/C.

Altogether, our results indicate that CycB3 promotes anaphase by acting upstream of the APC/C, not only in meiosis, but also in early embryonic mitoses, in collaboration with PP2A-Tws and independently from the SAC.

### CycB3 is required for correct anaphase and full APC/C activity in cells in culture

The above results indicate that maternally contributed CycB3 promotes mitotic anaphase in the early embryonic cell cycle. This is consistent with the observation that RNAi silencing of CycB3 delays mitotic anaphase in syncytial embryos (Yuan and O’Farrell, 2015). Since *CycB3* mutants were found to be viable and develop normally until adulthood, any potential role of CycB3 during mitosis in proliferating cells may have been overlooked. By contrast, although *CycB* mutants are also viable, roles have been reported for CycB in *Drosophila* cell division (Knoblich and Lehner, 1993; Wang and Lin, 2005). Moreover, CycB3 is expressed specifically in proliferating cells, and *CycB3* is required for viability when *CycB* is mutated, with double mutants displaying strong mitotic defects (Jacobs et al., 1998). To examine mechanistically how CycB3 contributes to mitotic regulation during divisions of proliferating cells, we used cells in culture. We generated D-Mel (d.mel-2) cells expressing H2A-RFP and Lamin-GFP, allowing us to measure the time between Nuclear Envelope Breakdown (NEB, T_0_) and anaphase onset. To silence CycB3, we tested 4 different dsRNAs, and chose dsRNA no 1 and 2 because they resulted in the most efficient depletion of CycB3 (Fig S2A). We found that silencing CycB3 delays anaphase entry (Fig 2A, Movies S1, S2). This is observed by a delay in the cumulative percentage of cells entering anaphase after NEB, with both CycB3 dsRNA 1 or 2 (Fig 2B, S2B). In addition, while control RNAi cells all enter anaphase within 3 hrs after NEB, a fraction of the CycB3-depleted cells were still arrested in metaphase after 3 hrs.

**Figure 2.**
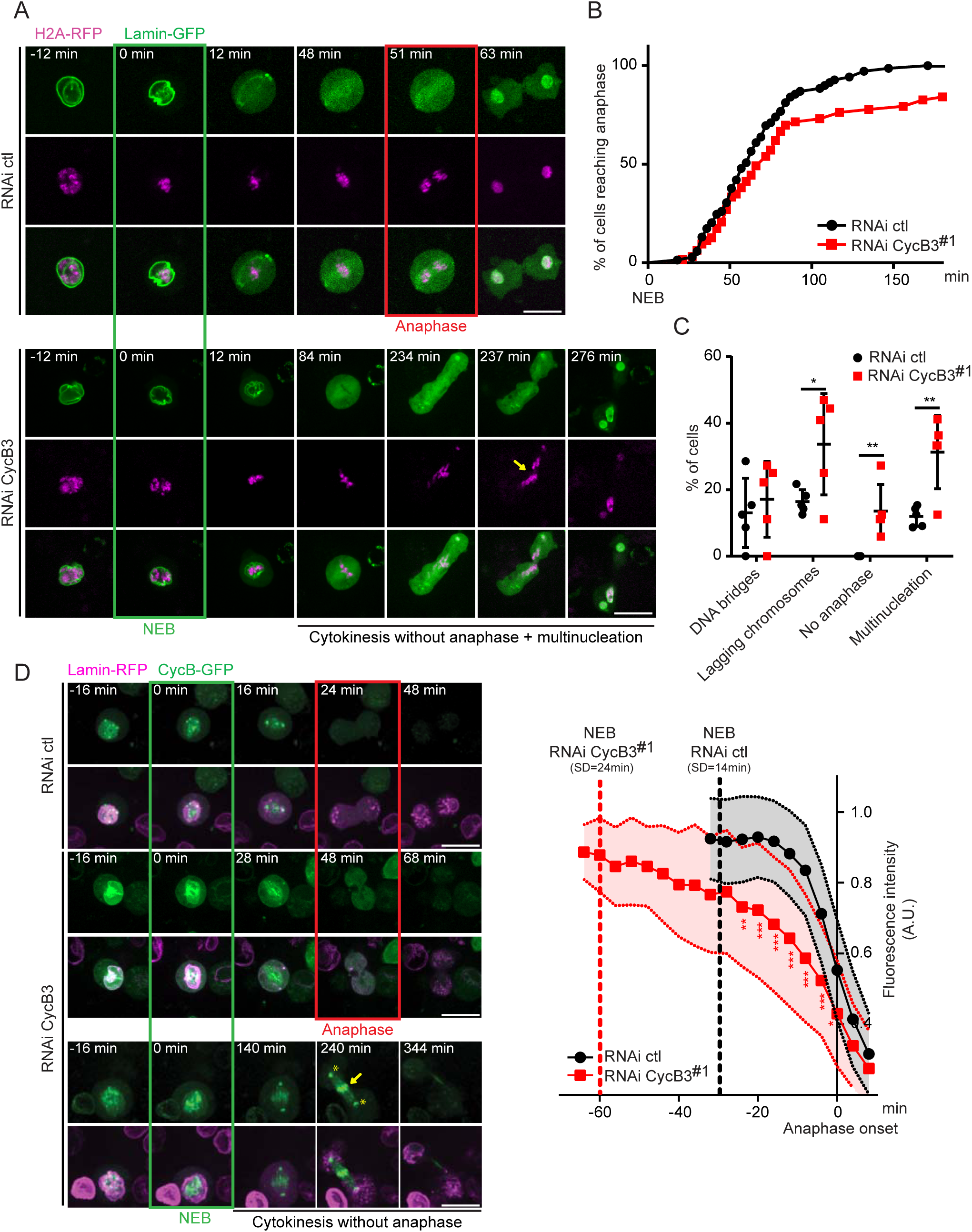
CycB3 is required for normal metaphase-anaphase transition and for CycB degradation in cells in culture. A. RNAi depletion of CycB3 delays anaphase entry and results in chromosome segregation defects in D-Mel cells. Note the cut through unsegregated chromosomes in this no-anaphase phenotype (arrow). RNAi ctl = transfection of dsRNA made from the bacterial KAN gene. B. Cumulative proportion of mitotic cells entering anaphase as a function of time after NEB. C. Frequency of cell division defects observed. Error bars: SD. For B-C, 63 and 69 cells were analyzed for RNAi CycB3 and ctl, respectively. D. RNAi depletion of CycB3 delays CycB degradation at the metaphase-anaphase transition. Left: examples of cell divisions (Z-projection of 3 planes in focus through the nucleus). Note the abnormal persistence of CycB-GFP on the chromosomes (arrow) and centrosomes (*) during cytokinesis. Right: levels of CycB-GFP quantified through time, relative to anaphase onset. 29 and 30 cells were analyzed for RNAi CycB3 and ctl, respectively. Error areas: SD. \*\*\**p* < 0.001; \*\**p* < 0.01; \**p* < 0.05 from paired *t*-tests. Scale bars: 10 µm. See also Figure S2.

Of the CycB3-depleted cells that do enter anaphase, a large proportion develop defects (Fig 2C, S2C). The frequencies of lagging chromosomes and post-mitotic multinucleation/micronucleation approximately double relative to control cells. Interestingly, CycB3-depleted cells that do not enter anaphase sometimes undergo cytokinesis, cutting through unsegregated chromosomes (Fig 2A, arrow, Movie S2). This is consistent with the previously observed role of CycB3 in negatively regulating cytokinetic furrowing (Echard and O’Farrell, 2003).

To determine if the anaphase delay and defects resulting from CycB3 silencing correlate with reduced APC/C activity, we monitored the levels of CycB, an essential target of the APC/C during mitotic exit. Our previous results showed that CycB levels are regulated by CycB3 in eggs (Fig 1D, 1F, S1A and S1B). We generated cells expressing CycB-GFP and Lamin-RFP, and measured CycB-GFP levels during mitosis. CycB-GFP is nuclear in prophase and becomes enriched on centrosomes and kinetochores until metaphase. We found that when CycB3 is depleted, CycB-GFP is degraded more slowly during the metaphase-anaphase transition, compared to control cells (Fig 2D, Movies S3, S4). In both conditions, anaphase onset occurs in the presence of similar CycB-GFP levels, suggesting that APC/C activity is rate-limiting for anaphase onset when CycB3 is depleted. Very similar results are obtained with a second dsRNA against CycB3 (Fig S2D). Interestingly, while CycB-GFP is not detected in control cells undergoing cytokinesis, it persists on centrosomes and chromosomes in CycB3-depleted cells that undergo cytokinesis without anaphase (Fig 2D and Movie S5).

We conclude that CycB3-dependent activation of the APC/C at the metaphase-anaphase transition is not restricted to female meiosis and the following syncytial embryonic mitotic cell cycles, but that it can also promote correct chromosome segregation during cell proliferation.

### CycB3 associates with the APC/C and is spatially regulated during the cell cycle

The above results demonstrate that CycB3 promotes APC/C activation. To begin exploring if this regulation is direct, we tested if CycB3 associates with the APC/C. We used D-Mel cells expressing tagged proteins to perform co-purifications experiments. We found that CycB3-PrA specifically co-purifies Myc-APC3 (Fig 3A). Very similar results are obtained with Myc-APC2 (Fig S3A). An interaction between CycB3 and the APC/C could occur as CycB3-Cdk1 binds the APC/C to phosphorylate it, or as CycB3 is recognized through its destruction box for ubiquitination by the APC/C. To discriminate between these modes of interaction, we used the destruction box mutation in CycB3. CycB3^D^-PrA is still able to co-purify Myc-APC3 or Myc-APC2 (Fig 3A, S3A), suggesting that the association observed does not reflect CycB3 binding as a substrate by the APC/C. We also found that CycB3-Myc can co-purify GFP-APC3 and GFP-APC6 in syncytial embryos (Fig S3B-C). Thus, CycB3 can form a complex with the APC/C in D-Mel cells and in syncytial embryos, possibly to directly regulate the APC/C.

**Figure 3.**
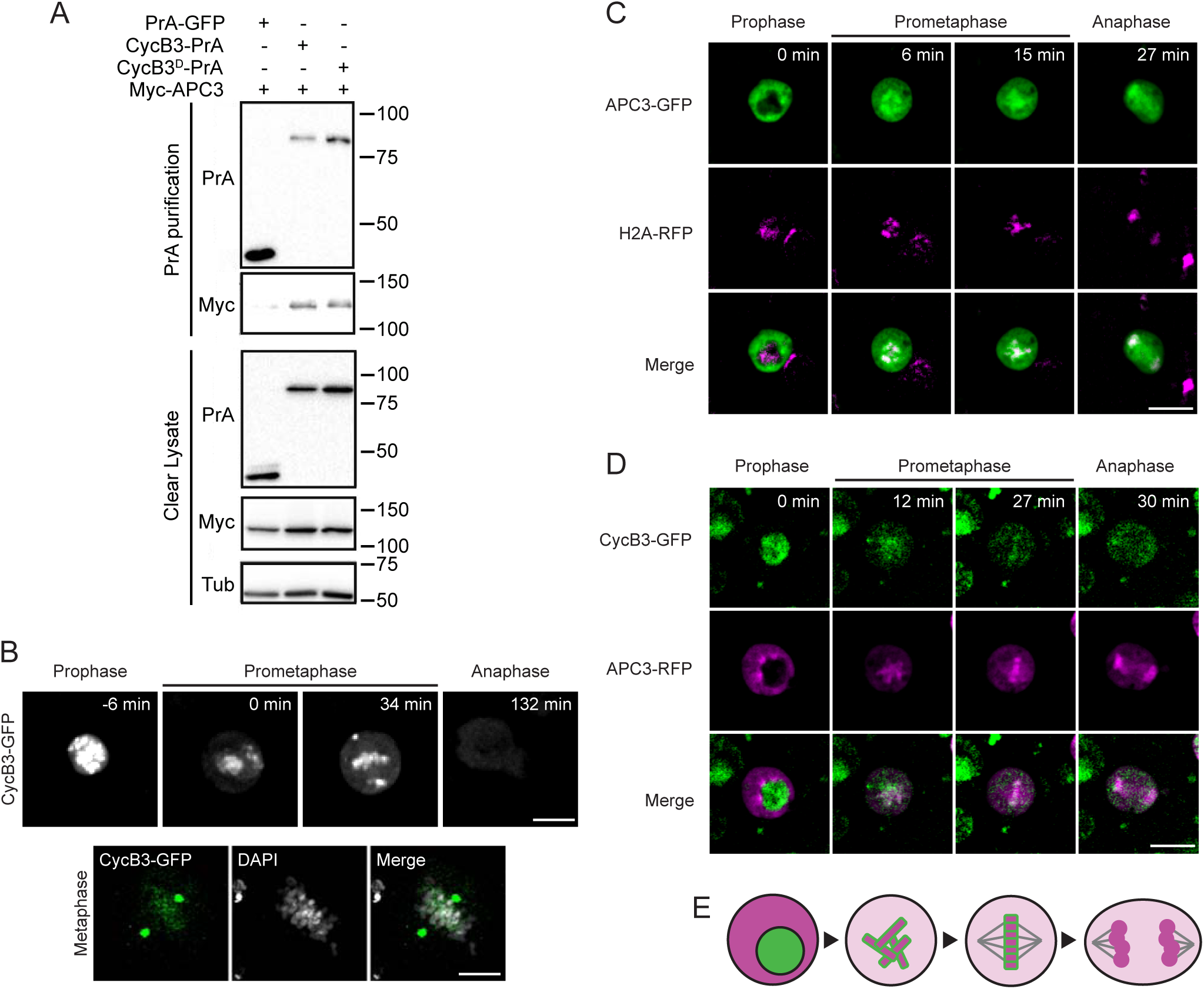
CycB3 associates with the APC/C and shows a dynamic localization relative to the APC/C in the cell cycle. A. Myc-APC3 associates with CycB3-PrA independently from its destruction box. Cells expressing the indicated proteins were submitted to Protein A affinity purification and products were analyzed by Western blot. B. CycB3-GFP localizes to chromosomes and spindle poles in early mitosis. Top: Live imaging; Bottom: a fixed cell in metaphase. C. APC3-GFP is predominantly cytoplasmic in interphase and becomes enriched on chromosomes during mitosis. D. CycB3-GFP and APC3-RFP are sequestered from each other in interphase and come together on chromosomes in mitosis. Scale bars: 10 µm except for the fixed cell: 5 µm. E. Schematic model of the dynamic localization of CycB3 (green) and the APC/C (magenta) from interphase to anaphase. See also Figure S3.

If CycB3 binds and directly activates the APC/C for the metaphase-anaphase transition, CycB3 and the APC/C should overlap in localization before anaphase. We investigated the spatio-temporal regulation of CycB3 relative to the APC/C using D-Mel cells. CycB3 overexpression appears to be toxic as it was difficult to obtain stable cell lines expressing CycB3 fused to any tag. Cells expressing the highest levels of CycB3-GFP tended not to divide. Nevertheless, we managed to catch divisions of cells expressing enough CycB3-GFP to reveal its dynamic localization. CycB3-GFP is strongly nuclear in interphase until prophase and becomes enriched on condensed chromosomes and on centrosomes as cells enter mitosis and until metaphase (Fig 3B, Movie S6). Starting in anaphase, CycB3-GFP levels are rapidly reduced, consistent with its degradation by the APC/C. Contrary to CycB3-GFP which is nuclear, APC3-GFP is strongly cytoplasmic in interphase until prophase. After NEB, APC3-GFP quickly becomes enriched on condensed chromosomes (marked by H2A-RFP) until telophase (Fig 3C), consistent with previous findings (Huang and Raff, 2002). We were able to generate a stable cell line where a fraction of the cells co-express detectable levels of CycB3-GFP and APC3-RFP. Examination of these cells confirms that nuclear CycB3-GFP and cytoplasmic APC3-RFP are mutually sequestered in interphase until NEB, and subsequently become co-enriched on or around chromosomes (Fig 3D-E, Movie S7).

Altogether, these results suggest that CycB3 interacts with the APC/C on chromosomes, to promote APC/C activation before anaphase.

### CycB3 promotes APC/C phosphorylation at CDK motifs and interaction with Cdc20 co-activators

The only known molecular function of *Drosophila* CycB3 is to bind and activate Cdk1. Thus, we next sought to determine if CycB3, in complex with Cdk1, promotes the activation of the APC/C by direct phosphorylation. To this end, we used syncytial embryos, where mitoses occur approximately every 10 min, thereby facilitating the detection of mitotic complexes and mitotic phosphorylation. We used fly lines expressing APC3/Cdc27 and APC6/Cdc16 N-terminally fused with GFP and expressed under the polyubiquitin promoter (Huang and Raff, 2002). We collected early embryos from these flies and purified the APC/C using a GFP-affinity resin. The band patterns we obtained after purification of GFP-APC3 or GFP-APC6 are similar on a silver-stained gel, and different from those obtained after purification of GFP-tagged Nup107, a component of the nuclear pore complex (Fig 4A). Mass spectrometry analysis revealed the presence of most APC/C subunits in the GFP-APC/C purification products (data not shown but see below). Western blots detected endogenous CycB3 that co-purified specifically with GFP-tagged APC/C, compared with GFP-Nup107 used as a negative control (Fig 4B). Cdk1 was also enriched in the GFP-APC/C purification products. Therefore, endogenous CycB3 and Cdk1 associate with the APC/C in early embryos.

**Figure 4.**
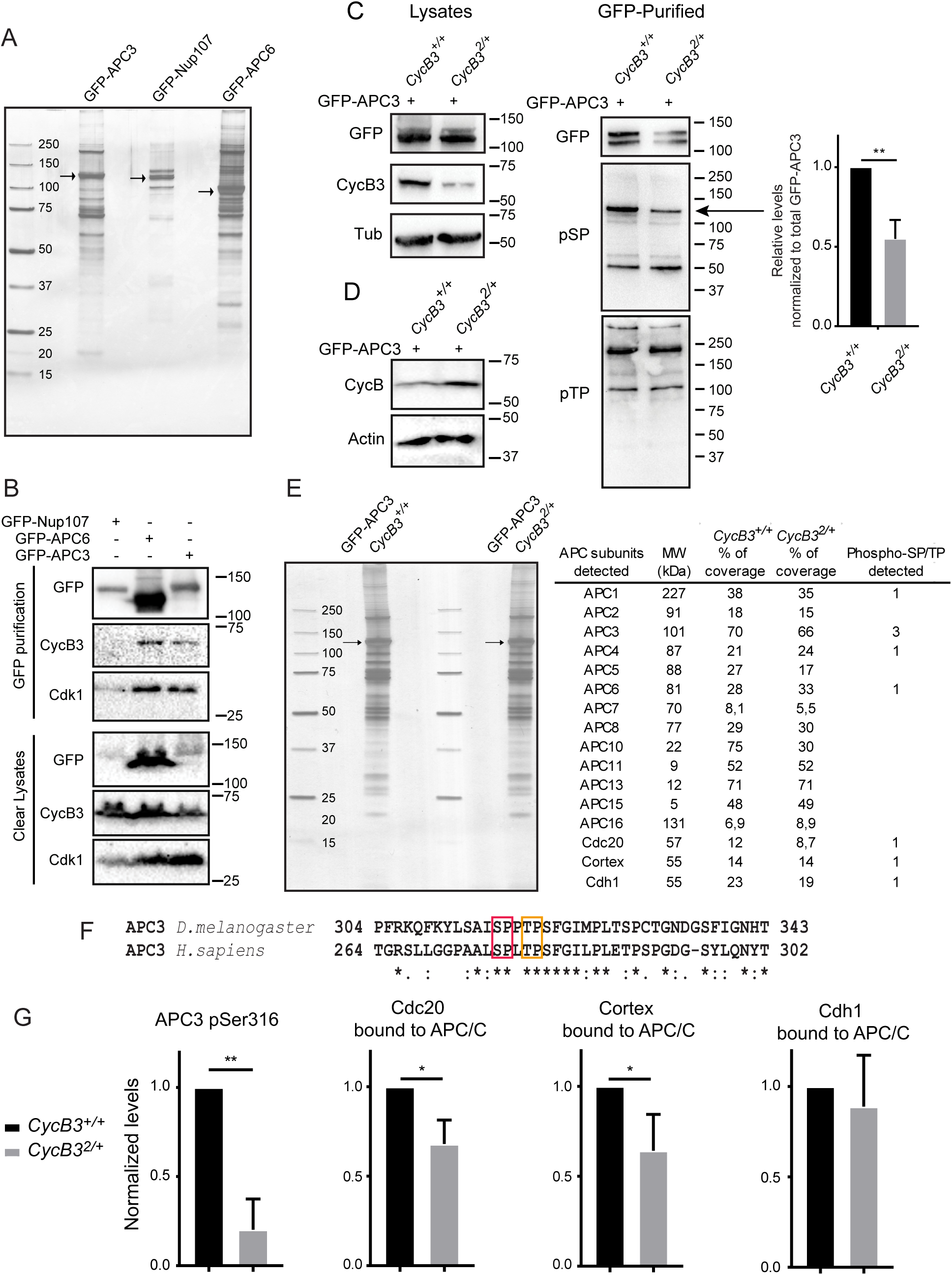
CycB3-Cdk1 physically associates with the APC/C and promotes its phosphorylation and interaction with Cdc20 co-activators. A. Embryos expressing GFP-APC3, GFP-APC6 and GFP-Nup107 were submitted to GFP-affinity purification and purified complexes were visualized on a silver-stained gel. Arrows: GFP-tagged proteins. B. CycB3 physically associates with the APC/C in embryos. Embryos expressing the indicated proteins were submitted to GFP-affinity purification and products were analyzed by Western blots. C. APC/C phosphorylation at SP site(s) is CycB3-dependent. GFP-affinity purifications were conducted from 0-2 hrs-old GFP-APC3 expressing embryos from *CycB3*^+*/*+^ and *CycB3*^*2/*+^ mothers and products were analyzed by Western blots. Left: Western blots on lysates used in the purifications. Note that CycB3 levels are lower in *CycB3*^*2/*+^ embryos; Center: GFP-APC3 purified products were analyzed by Western blot with antibodies against pSP or pTP sites (minimal CDK phosphorylation motifs). Arrow: presumed GFP-APC3 pSP band, less intense in *CycB3*^*2/*+^ embryos. Right: relative intensities of the band indicated by the arrow, after normalization to the amounts of total GFP-APC3 purified. Error bar: SD for 3 experiments. \*\**p* = 0.0082 from paired *t*-test. D. Western blot showing that CycB levels are higher in *CycB3*^*2/*+^ embryos (aged 0-2 hrs). E. Purification of the APC/C complex using GFP-APC3 from *CycB3*^+*/*+^ and *CycB3*^*2/*+^ embryos, for mass spectrometry analysis. Table: APC/C subunits detected by mass spectrometry. Their predicted molecular weight (MW), percentage of sequence detected and number of phospho SP/TP sites detected are indicated. F. Phosphorylation sites at minimal CDK motifs identified in *Drosophila* APC3 are conserved in the activation loop of human APC3. Red box: Ser316; Orange box: Thr319. G. Phosphorylation at Ser316 in APC3 is reduced in *CycB3*^*2/*+^ embryos. In addition, the amounts of Cdc20/Fizzy and Cortex co-purified with the APC/C are reduced in *CycB3*^*2/*+^ embryos. The amount of Cdh1/Fzr co-purified is not significantly affected. Proteins and phosphopeptides abundances were normalized to APC3 levels (see Methods). Error bars: SD for 3 experiments.

We then used phosphospecific Western blotting to explore whether the phosphorylation state of the APC/C depends on CycB3. For these experiments, we sought to compare the phosphorylation levels of purified APC/C from embryos laid by females wild-type for *CycB3* or heterozygous for the *CycB3*^*2*^ null allele. In embryos from *CycB3*^*2/*+^ mothers, the level of CycB3 is approximately halved, as expected (Fig 4C). Our genetic results indicating that a single copy of the *CycB3*^*2*^ allele is maternal-effect synthetic lethal with metaphase arrests in the context of heterozygosity for *tws* (Fig 1A-B) suggested that APC/C regulation may be significantly affected by this reduction in CycB3 levels. Consistent with this idea, CycB levels are higher in embryos from *CycB3*^*2/*+^ mothers (Fig 4D). We then purified the APC/C via GFP-APC3 and submitted the products to Western blots using phosphospecific antibodies against pSP or pTP sites, which are the minimal motifs for CDK phosphorylation. The anti-pSP blot reveals several bands (Fig 4C, center). After quantifications that included a correction to take into account slight variations in amounts of GFP-APC3 purified, we found only one pSP band that is significantly less intense in purified products from *CycB3*^*2/*+^ embryos (Fig 4C, arrow, quantified on right). This pSP band is the most intense in purification products from *CycB3*^+*/*+^ embryos, and its molecular mass of 125 kDa matches that observed for GFP-APC3. This result suggests that CycB3 is required for APC3 phosphorylation at CDK sites.

We set out to test this hypothesis directly using quantitative mass spectrometry. We repeated the purification of GFP-APC3 in both genotypes on a larger scale. Very similar amounts of APC/C were purified from *CycB3*^*2/*+^ or *CycB3*^+*/*+^ embryos, with similar band profiles detected on a gel (Fig 4E). We detected 3 phosphorylated sites at minimal CDK motifs in APC3, including S316 and T319. Both sites are conserved in human APC3, and map within the loop whose phosphorylation promotes APC/C activation. Moreover, both sites in human APC3 (S276 and T279) were shown to be phosphorylated *in vivo*, and by Cyclin A3-Cdk2 *in vitro* (Fig 4F) (Hegemann et al., 2011; Zhang et al., 2016). After normalization to total APC3 levels, we found that phosphorylation at S316 is strongly reduced in *CycB3*^*2/*+^ embryos vs *CycB3*^+*/*+^ embryos, suggesting that it is regulated by CycB3-Cdk1 (Fig 4G). Phosphorylation at T319 was too weak to be quantified by the same method. In addition, the Western blot with the anti-pTP reveals no obvious difference between *CycB3*^*2/*+^ and *CycB3*^+*/*+^ embryos (Fig 4C), suggesting that pTP sites on the APC/C are not major points of regulation by CycB3. The third site we identified, S455, does not change in intensity of phosphorylation between genotypes (not shown).

CDK phosphorylation of APC3 is an essential part of a mechanism for APC/C activation demonstrated *in vitro* (Fujimitsu et al., 2016; Zhang et al., 2016). This event primes the APC/C for its CDK phosphorylation of APC1, which results in the displacement of a switch loop within APC1, opening the binding site for Cdc20. There are three Cdc20 paralogs in *Drosophila*: Fizzy/Cdc20 that activates the APC/C in mitosis and meiosis (Dawson et al., 1995; Pesin and Orr-Weaver, 2007; Sigrist et al., 1995; Swan and Schupbach, 2007), Cortex that activates the APC/C in female meiosis and possible syncytial mitoses (Pesin and Orr-Weaver, 2007; Swan and Schupbach, 2007), and Cdh1 which is not detected in syncytial embryos and keeps the APC/C active in G1 in later developmental stages (Raff et al., 2002). To test the idea that CycB3 promotes APC binding to its co-activators, we determined the relative amounts of Cdc20, Cortex and Cdh1 in GFP-APC3 purification products from *CycB3*^*2/*+^ and *CycB3*^+*/*+^ embryos. Cdc20, Cortex and Cdh1 are all detected in association with the APC/C purified from embryos aged from 0-2 hrs (Fig 4E). The presence of Cdh1 may be surprising but it could come from the oldest embryos collected that may have initiated the maternal-zygotic transition. Strikingly, the relative levels of both Cdc20 and Cortex are lower in the GFP-APC3 purification products from *CycB3*^*2/*+^ embryos compared with *CycB3*^+*/*+^ embryos (Fig 4G). However, Cdh1 levels are unchanged. These results suggest that CycB3 specifically promotes APC/C recruitment of its mitotic and meiotic co-activators Cdc20 and Cortex.

### Mechanism and conservation of the role of Cyclin B3 in the activation of the APC

Altogether, our results strongly suggest that CycB3-Cdk1 directly activates the APC/C by phosphorylation, promoting its function at the metaphase-anaphase transition in meiosis and in both maternally driven early embryonic mitoses and somatic cell divisions. This regulation is likely mediated by the phosphorylation in the activation loop of APC3 by CycB3-Cdk1 that ultimately promotes the recruitment of the Cdc20-type co-activators Fizzy and Cortex (Fig 5). Previous work has shown that APC3 phosphorylation and APC/C activation by cyclin-CDK complexes require their CKS subunit (Patra and Dunphy, 1998; Shteinberg and Hershko, 1999; Zhang et al., 2016). CKS subunits can act as processivity factors that bind phosphorylated sites to promote additional phosphorylation by the CDK (Bourne et al., 1996; Koivomagi et al., 2011). Thus, phosphorylation of APC3 would prime the binding of a cyclin-CDK-CKS complex, to promote the additional phosphorylation of APC1, allowing for Cdc20 binding (Zhang et al., 2016). The mutation of phosphorylation sites into Asp or Glu residues cannot substitute for the presence of phosphate in the CKS binding site, precluding the use of phosphomimetic mutations to test the model (McGrath et al., 2013). It is likely that our analysis did not detect all phosphorylation sites in the APC/C. Thus, we cannot exclude the possibility that other phosphorylation events, mediated by CycB3-Cdk1 or another kinase, may be required for complete APC/C activation. For example, other phosphorylation events have been proposed to regulate APC/C localization (Huang et al., 2007). The interdependence between CycB3 and Tws that we uncovered may reflect a role of PP2A-Tws in the recruitment of Cdc20 co-activators to the APC/C. Cdc20 must be dephosphorylated at CDK sites before binding the APC/C, and in human cells both PP2A-B55 and PP2A-B56 promote this event (Fujimitsu and Yamano, 2020; Hein et al., 2017; Labit et al., 2012).

**Figure 5.**
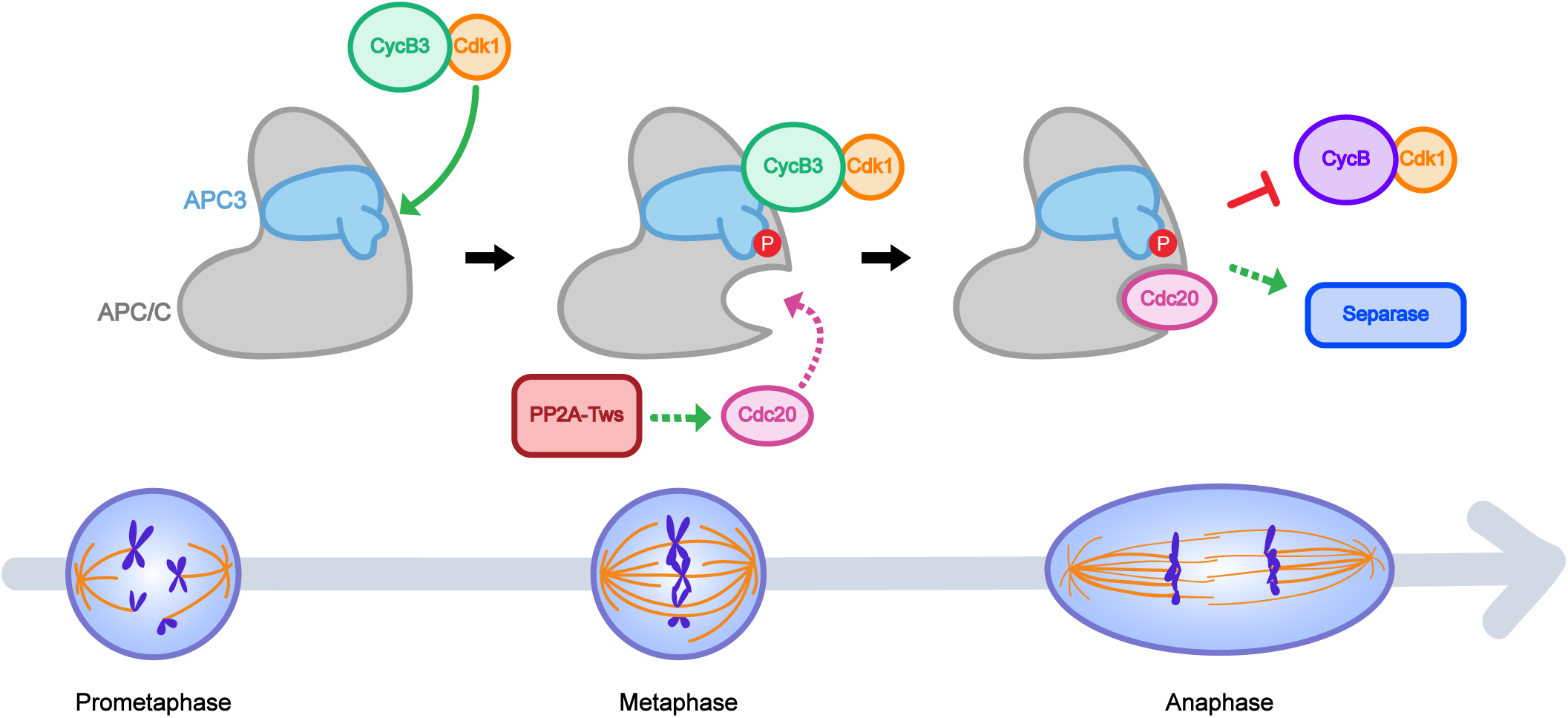
Model for APC/C activation by CycB3-Cdk1 in meiosis and mitosis. CycB3-Cdk1 binds the APC/C and phosphorylates its APC3 subunit. This event promotes the binding of Cdc20 co-activators, likely through known mechanisms (Fujimitsu et al., 2016; Zhang et al., 2016). In addition, PP2A-Tws may help Cdc20 recruitment to the APC/C (Hein et al., 2017; Labit et al., 2012). Activated APC/C triggers CycB degradation, separase activation and anaphase.

CycB3 is strongly required for APC/C activation in meiosis and in the early syncytial mitoses, and to a lesser extent in other mitotic divisions, despite the presence of two additional mitotic cyclins, CycA and CycB, capable of activating Cdk1. There are many possible reasons for this requirement. Overexpression of stabilized forms of CycA or CycB can block or slow down anaphase, suggesting that they may interfere with APC/C function in this transition (Parry and O’Farrell, 2001). However, under normal expression levels, CycA or CycB or both may contribute to activate the APC/C like CycB3. *CycB3* mutant flies develop until adulthood (Jacobs et al., 1998), which implies that the APC/C can be activated to induce anaphase in at least a vast proportion of mitotic cells, and this activation could be mediated by CycA and/or CycB. *CycA* is essential for viability and *CycB* mutants show strong female germline development defects, complicating the examination of potential roles for these cyclins at the metaphase-anaphase transition (Bourouh et al., 2016; Jacobs et al., 1998; Lehner and O’Farrell, 1989; Wang and Lin, 2005). Thus, in principle, the requirements for CycB3 in female meiosis, in embryos and in mitotic cells in culture could merely reflect the need for a minimal threshold of total mitotic cyclins. We consider this possibility unlikely because CycB3 is expressed at much lower levels than CycB in early embryos (Casas-Vila et al., 2017). Moreover, while maternal heterozygosity for mutations in *CycB3* and *tws* causes a metaphase arrest in embryos (Fig 1), heterozygosity for mutations in *CycB* and *tws* does not cause embryonic defects (Mehsen et al., 2018). In fact, genetic results suggest that the function of CycB is antagonized by PP2A-Tws in embryos, while CycB3 and PP2A-Tws collaborate for APC/C activation in embryos (Mehsen et al., 2018). Thus, although it is possible that CycA and CycB can participate in APC/C activation, CycB3 probably has some unique feature that makes it particularly capable of promoting APC/C activation.

By what mechanism could CycB3 be particularly suited for APC/C activation? Cyclins can play specific roles by contributing to CDK substrate recognition or by directing CDK activity in space and time (Bloom and Cross, 2007). We did not investigate the precise nature of the molecular recognition of the APC/C by CycB3. It may be that CycB3 possesses a specific binding site for the APC/C that is lacking in CycA and CycB. Another possibility is that differences in localization between cyclins dictate their requirements. In particular, while CycA and CycB are cytoplasmic in interphase, CycB3 is nuclear (Jacobs et al., 1998). We surmise that the nuclear localization of CycB3 may help concentrate CycB3 in the spindle area upon germinal vesicle breakdown, when the very large oocyte enters meiosis. In future studies, it will be interesting to compare the ability of different mitotic cyclins to activate the APC/C and to determine the molecular basis of potential differences.

In any case, our results show that CycB3 activates the APC/C and that this regulation is essential in *Drosophila*. Cyclin B3 has been shown to be required for anaphase in female meiosis of insects (*Drosophila*), worms (*C. elegans*) and vertebrates (mice). It is tempting to conclude that the activation of the APC/C is a function of Cyclin B3 conserved in all these species. In fact, expression of Cyclin B3 from *Drosophila*, zebrafish or *Xenopus* can rescue female meiotic defects in *Ccnb3* knock-out mice (Karasu et al., 2019). In mouse oocytes, like in flies, the metaphase arrest that results from Cyclin B3 inactivation was shown to be independent from the SAC (Karasu et al., 2019; Li et al., 2019b; Zhang et al., 2015a), consistent with a direct activation of the APC/C by CycB3. However, in *C. elegans* embryos, the metaphase arrest upon CYB-3 (Cyclin B3) inactivation requires SAC activity (Deyter et al., 2010). The underlying mechanism and whether it also occurs in other systems remain to be determined. However, CYB-3 plays roles in *C. elegans* that have not been detected for Cyclin B3 in flies or vertebrates, including a major role in mitotic entry, where CYB-3 mediates the inhibitory phosphorylation of Cdc20 (Lara-Gonzalez et al., 2019). In this regard, *C. elegans* CYB-3 may be more orthologous to Cyclin A. Yet, given that Cyclin B3 is required for anaphase in a SAC-independent manner in flies and mice, it seems reasonable to suggest that the direct activation of the APC/C by Cyclin B3 is conserved in vertebrates.

## MATERIALS AND METHODS

### Plasmid and mutagenesis

Plasmids were generated using the Gateway recombination system (Invitrogen). Coding sequences were first cloned into a pDONOR221 entry vector. Then, they were recombined into the relevant destination vectors for expression under either the copper-inducible (pMT) or constitutive (pAC5) promoters. The following expression vectors were generated: pAC5-CycB-GFP, pAC5-CycB3-GFP, pMT-CycB3-GFP, pAC5-PrA-GFP, pAC5-CycB3-PrA, pAC5-CycB3^D^-PrA, pAC5-Myc-APC3, pAC5-Myc-APC2, pMT-APC3-GFP, pMT-APC3-RFP, pMT-H2A-RFP, pMT-Lamin-RFP and pAC5-Lamin-GFP. For CycB3^D^, the mutagenesis was performed using *PFU* DNA Polymerase (Biobasic) according to the manufacturer’s instructions.

### Fly genetics

Fly culture was done according to standard procedures. All crosses were performed at 25°C. The WT strains used were *W*^*1118*^ and *Oregon R*. Transgenic flies for the expression of pUASp-CycB3-Myc and pUASp-GFP-CycB3^D^ were generated by random insertion.

The Bloomington *Drosophila* Stock Center (BDSC) provided the following fly lines for RNAi: White^GL00094^ (35573), APC1^HMS01744^ (38531), APC3^HMC03814^ (55155); for mutant alleles: *CycB3*^*2*^ (6635), *CycB3*^*L6*^ (*CycB3*^*L6540*^; 10337), *mad2*^*EY21687*^ (22495), *cort*^*QW55*^(4974), *cort*^*RH65*^(77860) and *C(1)M3, y*^*2*^*/C(1;Y)1, y*^*1*^*/0* (38416); for transgenic insertions: *GFP-Nup107* (35514), and for drivers: *matα4-GAL-VP16* (7062) and *otu-Gal4-VP16* (58424). Fly strains harboring the *CycB3*^*2*^ and *CycB3*^*L6*^ alleles and the GFP-Nup107 insertion were from the BDSC. The *tws*^*P*^ and *tws*^*aar1*^ alleles were obtained from David Glover. The *fzy*^*6*^ and *fzy*^*7*^ alleles were obtained from Ian Dawson. pUb-GFP-APC3 and pUb-GFP-APC6 lines were provided by Yuu Kimata and were previously described (Huang and Raff, 2002).

To generate *cort*^*QW55/RH6*^, *fzy*^*6/7*^ double mutant females, *cort*^*QW55*^ was recombined with *fzy*^*6*^ and *cort*^*RH65*^ was recombined with *fzy*^*7*^. Crosses to generate *cort*^*QW55/RH65*^, *fzy*^*6/7*^ double mutants were reared at 18°C until pupation and then moved to 29°C. Wild type unfertilized eggs were generated by crossing wild type females to X/O males. X/O males were generated by crossing *X^Y (C(1)M3, y*^*2*^*/C(1;Y)1, y*^*1*^*/0*) males to wild type females.

Expression of *UASp* transgenes in the female germline and early embryo were done with *matα4-GAL-VP16* in Figure 1A, C and S1A, and with *otu-Gal4-VP16* in Figures S1B.

For fertility tests, between 5 and 10 individual females were crossed with 3 males and allowed to lay eggs for 1 day on grape juice agar tubes with yeast paste before being removed. The percentage of hatched eggs was counted 24 h later.

### *Drosophila* cell culture and transfections

All cells were in the D-Mel (d.mel-2) background and were cultured in Express Five medium supplemented with glutamine, penicillin, and streptomycin. All stable cell lines were selected in medium containing 20 µg/ml blasticidin. Expression of the pMT constructs was induced with 300 μM CuS0_4_ overnight.

Plasmid transfections were performed using X-tremeGENE HP DNA Transfection Reagent (Roche). For RNA interference, cells were transfected using Transfast reagent in six-well plates with 30 μg of dsRNA against CycB3 or control. The control dsRNA was generated against the sequence of the bacterial kanamycin resistance gene (*KAN*). Cells were analyzed 48 h later by immunoblotting or live-cell imaging.

### Protein purifications

Protein A affinity purifications from D-Mel cells were done largely as described (D’Avino et al., 2009). Briefly, cells were co-transfectd with pAC5-CycB3-Pra and pAC5-Myc-APC2 or pAC5-Myc-APC3, harvested from confluent 75 cm^2^ flasks and resuspended in 1.6 ml of lysis buffer (50 mM Tris-CL pH 7.5, 150 mM NaCl, 0.1 mM EGTA, 2 mM MgCl_2_, 1 mM DTT, 1.5 µM aprotinin, 23 µM leupeptin, 1 mM PMSF, 10% glycerol, 0.5% triton X-100) supplemented with protease inhibitors. Lysates were clarified by centrifugation at 10 000 g for 15 min in a tabletop centrifuge at 4 °C. Supernatants were incubated for 1 h at 4 °C with 25 μl of DynaBeads previously conjugated to rabbit IgG. Beads were washed five times with 1 ml of lysis buffer for 5 min at 4°C. Purification products were eluted by heating at 95 °C for 5 min in Laemmli buffer and analyzed by Western blotting.

For protein immunopurifications from egg/embryos, 0-2 h collections were performed at 25°C. Eggs were dechorionated in 50% bleach, then washed in PBS and frozen in liquid nitrogen. 100 mg of embryos were crushed in lysis buffer as above. Lysates were incubated for 30 min at 4°C on a rotating wheel, and centrifugated at 10 000 g for 15 min. Lysates were then incubated with antibody (anti-Myc antibody 9E10 or anti-CycB3, from rabbit, custom-made by Genscript) for 1 hour. Then the lysate was incubated for 2 hours with PrG-coupled (for anti-Myc) or PrA-coupled magnetic beads (anti-CycB3). Beads were washed 5 times in lysis buffer followed by elution in Laemmli buffer at 95°C for 5 min.

For GFP affinity purifications, embryos were crushed in the same lysis buffer as above but supplemented with 1% phosphatase inhibitor (Cocktail 2 (P5726) and 3 (P0044), Sigma). Lysates were incubated for 30 min at 4°C on a wheel, and centrifugated at 10 000 g for 15 min. Lysates were then incubated with GFP-trap nanobeads (Chromotek) for 2 h. Beads were washed 5 times with lysis buffer. For mass spectrometry, 5 additional washes in PBS + protease inhibitors were performed. 90% of the sample was sent for mass spectrometry. The remaining 10% was eluted in Laemmli buffer and used for SDS-PAGE analysis by silver nitrate staining and Western blots.

### Mass spectrometry

Samples were reconstituted in 50 mM ammonium bicarbonate with 10 mM TCEP [Tris(2-carboxyethyl)phosphine hydrochloride; Thermo Fisher Scientific], and vortexed for 1 h at 37°C. Chloroacetamide (Sigma-Aldrich) was added for alkylation to a final concentration of 55 mM. Samples were vortexed for another hour at 37°C. One microgram of trypsin was added, and digestion was performed for 8 h at 37°C. Supernatants were desalted on stage-tips (The Nest Group). Samples were dried down and solubilized in 5% ACN-0.2% formic acid (FA). The samples were loaded on a home-made reversed-phase column (150-μm i.d. by 150 mm) with a 220-min gradient from 10 to 30% ACN-0.2% FA and a 600-nl/min flow rate on a Easy nLC-1000 connected to an Orbitrap Fusion (Thermo Fisher Scientific, San Jose, CA). Each full MS spectrum acquired at a resolution of 240,000 was followed by tandem-MS (MS-MS) spectra acquisition on the most abundant multiply charged precursor ions for a maximum of 3s. Tandem-MS experiments were performed using collision-induced dissociation (CID) at a collision energy of 30%. The data were processed using PEAKS X (Bioinformatics Solutions, Waterloo, ON) and a Uniprot *Drosophila* unreviewed database. Mass tolerances on precursor and fragment ions were 10 ppm and 0.3 Da, respectively. Fixed modification was carbamidomethyl (C). Variable selected posttranslational modifications were oxidation (M), deamidation (NQ), phosphorylation (STY), acetylation (N-ter).

The data were visualized with Scaffold 4.3.0. For measurements of relative abundances, areas under precursor peptide peaks from LC-MS spectra were obtained through PEAKS X quantitation mode. Normalization of peptide areas was performed from total ion current of all LC-MS acquisitions. The top 3 most intense peptide areas from the LC-MS were summed to give protein abundances.

### Western blotting and immunofluorescence

Primary antibodies used in Western blotting and immunofluorescence were anti-GFP from rabbit (TP401, Torrey Pines, at 1:2000 dilution for WB), anti-GFP from rabbit (A6455, Invitrogen, at 1:200 for IF), anti-CycA from mouse (A12 purified, Developmental Studies Hybridoma Bank, 1/1000), anti-CycB from mouse (F2F4 purified, Developmental Studies Hybridoma Bank, 1/2000), anti-CycB3 from rabbit (Custom-made by Thermo Fisher Scientific, at 1:2000 dilution for WB; except for Fig 1D, where a rabbit antibody provided by Christian Lehner was used), anti-B55 (tws) from rabbit (2290S, Cell Signaling, at 1:2000 for WB) anti-α-tubulin DM1A from mouse (#T6199, Sigma, at 1:10,000 dilution for WB), anti-Myc 9E10 from mouse (#sc-40, Santa Cruz Biotechnology, Inc., at 1:2000 dilution for WB), mouse monoclonal anti-actin (#MAB1501, Millipore, at 1/5000), anti-P-SP from rabbit (#9111, Cell Signaling, at 1:2000 dilution for WB in 5%BSA), anti-P-TP from rabbit (#14371, Cell Signaling, at 1:2000 dilution for WB in 5% BSA), anti-Cdk1 PSTAIR from mouse (#10345, Abcam, at 1:2000 dilution for WB), anti-α-tubulin YL1/2 from rat (MCA77G, Bio-Rad AbD Serotec, at 1:2000 dilution for IF), anti-rabbit Alexa Fluor 488 (A11008, Invitrogen, at 1/200 for IF), anti-rat Alexa Fluor 647 from goat (A21247, Invitrogen, at 1:1000 for IF), peroxidase-conjugated anti-mouse from goat (115-035-003, Jackson ImmunoResearch, at 1:5000 for WB), peroxidase-conjugated anti-rabbit from goat (111-035-008, Jackson ImmunoResearch, at 1:5000for WB), peroxidase-conjugated ChromPure from Rabbit (011-030-003, Jackson ImmunoResearch, at 1:5000 for WB).

For Western blots from stage 14 oocytes, ovaries were dissected in Isolation Buffer (55 mM NaOAc, 40 mM KOAc, 110 mM sucrose, 1.2 mM MgCl2, 1 mM CaCl2, and 100 mM HEPES, pH 7.4) with collagenase to dissociate individual oocytes. Late stage oocytes were selectively enriched by repeated rinses in Isolation Buffer and removing slowly settling smaller egg chambers.

### Microscopy

For immunofluorescence, cells were fixed on coverslips with 4% formaldehyde during 20 min at room temperature (RT). Cells were permeabilized and blocked in PBS containing 0.1% Triton X-100 and 1% BSA. Cells were incubated with primary antibodies diluted in PBS containing 0.1% Triton (PBT) for 1 h at RT, washed three times in PBS and incubated with secondary antibodies and DAPI diluted in PBT for 1 h at RT. Then, coverslips were washed 3 times in PBS before being mounted in Vectashield medium (Vector Laboratories). Imaging was performed using a confocal system Zeiss LSM880.

For Fluorescence *In Situ* Hybridization (FISH), 0-2 h eggs were dechorionated in bleach and then washed 3 times in 0.7% NaCl, 0.05% Triton X-100). Eggs were then fixed in methanol:heptane (1:1) while shaking vigorously. Eggs were then stored at −20°C for later use, or immediately rehydrated successively in 9:1, 7:3 and 1:1 methanol:PBS solutions. FISH was performed on methanol-fixed eggs as described (Dernburg, 2000), using a probe against the 359-base pair peri-centromeric repeat on the X chromosome. DNA was stained with QUANT-IT Oligreen (#O7582, Invitrogen, 1/5000). Eggs were mounted using tetrahydronaphthalene. Imaging was performed using a confocal system Leica SP8.

For live analysis of *Drosophila* syncytial embryos, 0-2 h embryos were first dechorionated in 50% bleach, aligned on a coverslip and covered with halocarbon oil. 25 confocal sections of 1 μm were collected per time point for each embryo. Live imaging was performed using a Spinning-Disk confocal system Yokogawa CSU-X1 5000 mounted on a fluorescence microscope (Zeiss Axio Observer.Z1). Images were treated using Zen software (Zeiss) and Fiji software (NIH) as described (Kachaner et al., 2017).

To film D-Mel cells, they were plated on a LabTek II chambered coverglass (#155409, Thermo Fisher Scientific) and imaging was performed using the same Spinning-Disk microscope as for embryos. To monitor CycB-GFP levels, GFP fluorescence signal in the whole cell was quantified directly with the Zen software. To measure the GFP fluorescence ratio, time-lapse images were collected at 3 min intervals for H2A-RFP Lamin-GFP cells or 4 min intervals for Lamin-RFP CyclinB-GFP cells. For each timepoint, a circle was drawn in a single in-focus plane at the level of chromosomes and the whole intensity was measured. A ratio was then calculated between each timepoint value and the first timepoint value.

## ACKNOWLEDGEMENTS

We thank Myreille Larouche for making Figure 5, and David Kachaner and other members of the Archambault lab for useful discussions. We are grateful to Christian Charbonneau for help with the microscopy. We thank Christian Lehner for a gift of anti-Cyclin B3 antibody, and Yuu Kimata for pUb-GFP-APC3 and pUb-GFP-APC6 flies. This work was supported by Discovery grants from the Natural Sciences & Engineering Research Council of Canada to VA and AS, and by an Operating grant from the Cancer Research Society of Canada to VA. DG was a recipient of a postdoctoral fellowship from the Fonds de Recherche du Québec – Santé (FRQS). The authors declare no competing financial interests.

## AUTHOR CONTRIBUTIONS

DG, MB, EB, PT, AS and VA designed experiments and wrote the paper. DG, MB and EB performed experiments.

