## Supplementary Figures and Legends for "Cyclin B3 activates the Anaphase-Promoting Complex/Cyclosome in meiosis and mitosis"

**Figure S1. CycB3 promotes APC/C activity in eggs (related to Fig 1).** A-B. CycB3 negatively regulates CycB levels in an APC-dependent manner. Eggs were collected for 2 hrs from the indicated conditions and analyzed by Western blot. In A, expression of *UASp-GFP-CycB3*$^{D}$ and of *UASp-APC1 RNAi* was driven maternally by *Mat α-Tubulin Gal4-VP16*. The *UASp-GFP-CycB3*$^{D}$ and *UASp-APC1 RNAi* alone genotypes also contained a *UASp-WHITE* construction, to control for potential dilution of Gal4. Eggs from all conditions failed to develop (Ctl: unfertilized eggs). In B, expression of *UASp-GFP-CycB3*$^{D}$ was driven maternally by *otu-Gal4-VP16.* Error bars: SD. **$p < 0.01$; *$p < 0.05$ from paired $t$-tests.

**Figure S2. CycB3 promotes APC/C activity in mitosis in D-Mel cells (related to Fig 2).** A. dsRNAs targeting different regions of the CycB3 transcript were tested. No 1 and 2 were more effective and selected for experiments. B-D. Similar results were obtained with dsRNA no 2, compared with dsRNA no 1 (Fig 2). B-C. RNAi ctl: 106 cells; RNAi CycB3: 107 cells analyzed. D. RNAi ctl: 30 cells; RNAi CycB3: 40 cells analyzed. Error areas and error bars: SD. ***$p < 0.001$; **$p < 0.01$; *$p < 0.05$ from paired $t$-tests. Scale bars: 10 µm.

**Figure S3. CycB3 associates with the APC/C (related to Fig 3).** A. Myc-APC2 associates with CycB3-PrA independently from its destruction box. Cells expressing the indicated proteins were submitted to Protein A affinity purification and products were analyzed by Western blot. B-C. CycB3-Myc co-purifies GFP-APC3 (B) or GFP-APC6 (C) in *Drosophila* syncytial embryos.

**Movie S1. Normal mitotic division of a D-Mel cell expressing Lamin-GFP and H2A-RFP after transfection with the KAN dsRNA (control).** Z-projection of 3 plans in focus of the nucleus. Scale bars: 10 µm.

**Movie S2. Abnormal mitotic division of a D-Mel cell expressing Lamin-GFP and H2A-RFP after transfection with dsRNA against CycB3.** Anaphase is delayed and chromosome segregation is defective. Z-projection of 3 plans in focus of the nucleus. Scale bars: 10 µm.

**Movie S3. CycB is rapidly degraded before anaphase onset in a D-Mel cell.** Normal mitosis in a D-Mel cell expressing CycB-GFP and Lamin-RFP transfected with KAN dsRNA (control). CycB-GFP localizes on the spindle, kinetochores and centrosomes during mitosis. Z-projection of 3 plans in focus of the nucleus. Scale bars: 10 µm.

**Movie S4. CycB degradation is slower and anaphase is delayed in a dividing D-Mel cell depleted of CycB3.** Cell expressing CycB-GFP and Lamin-RFP and transfected with CycB3 dsRNA. Z-projection of 2 plans in focus of the nucleus. Scale bars: 10 µm.

**Movie S5. CycB degradation is incomplete in a D-Mel cell depleted of CycB3 undergoing cytokinesis without anaphase**. Cell expressing CycB-GFP and Lamin-RFP and transfected with CycB3 dsRNA. CycB-GFP persists on the chromosomes and centrosomes during cytokinesis. Z-projection of 3 plans in focus through the nucleus. Scale bars: 10 µm.

**Movie S6. CycB3 localization dynamics in a dividing D-Mel cell.** CycB3-GFP localizes to chromosomes and spindle poles in early mitosis. Z-projection of 3 plans in focus through the nucleus. Scale bars: 10 µm.

**Movie S7. Relative localizations of CycB3 and the APC/C in a dividing D-Mel cell.** CycB3-GFP and APC3-RFP both localize to chromosomes before anaphase. Z-projection of 3 plans in focus through the nucleus. Scale bars: 10 µm.

Figure S1

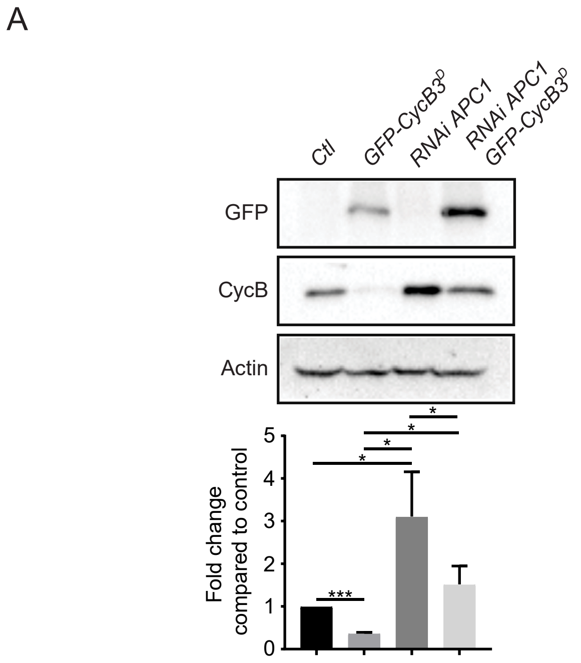
A
Ctl
GFP-CycB3D
RNAi APC1
RNAi APC1 GFP-CycB3D
GFP
CycB
Actin
Fold change compared to control
0
1
2
3
4
5
***
*
*
*
*

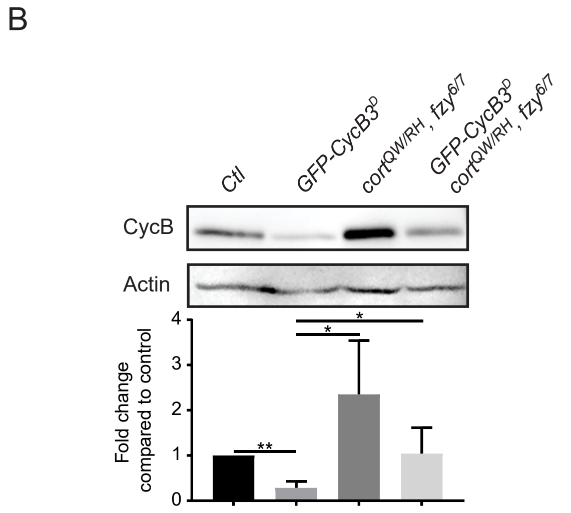
B
Ctl
GFP-CycB3D
cortQW/RH, fzy6/7
GFP-CycB3D cortQW/RH, fzy6/7
CycB
Actin
Fold change compared to control
0
1
2
3
4
**
*
*

### Figure S2

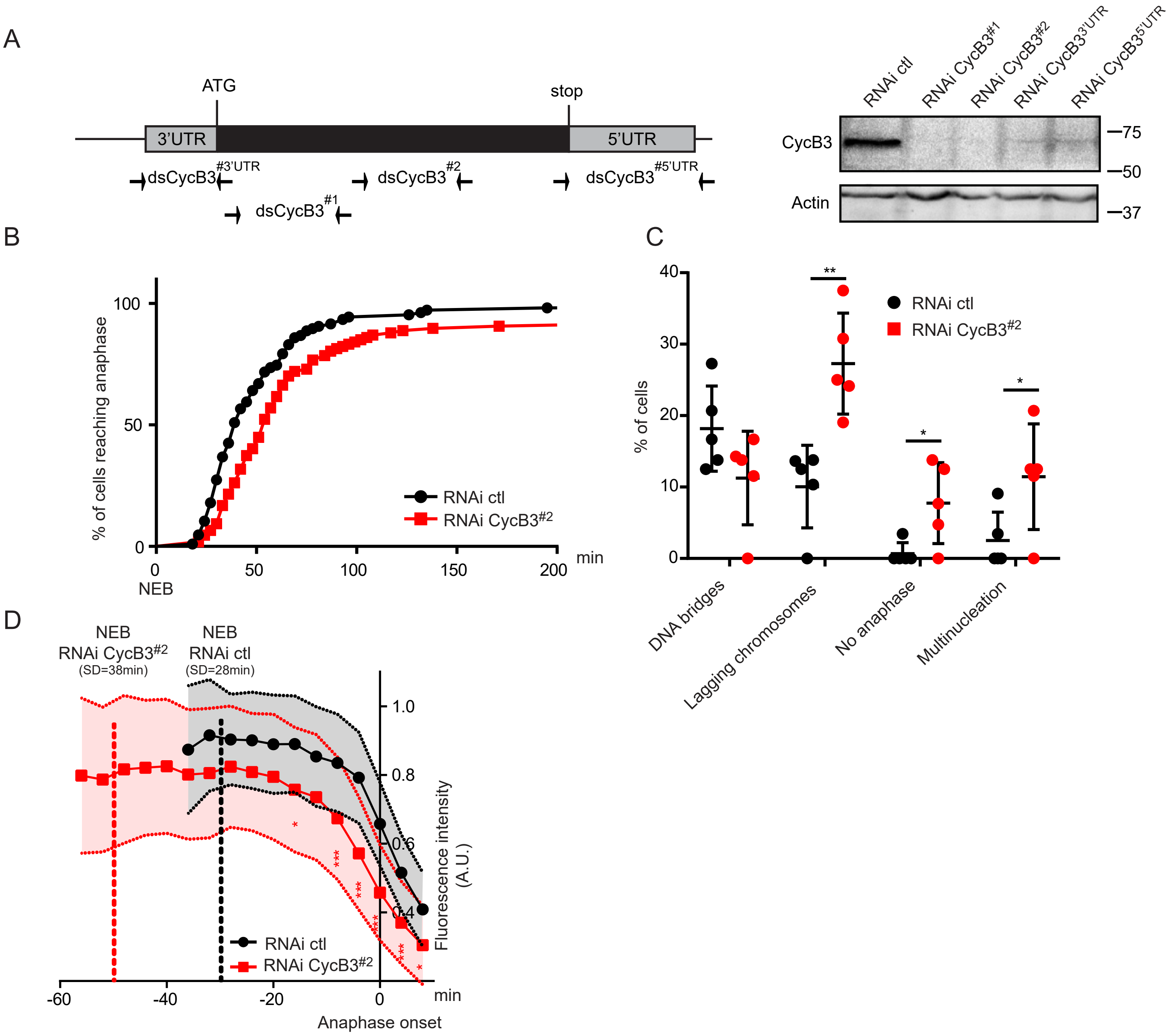
A
ATG
stop
3'UTR
5'UTR
dsCycB3#3'UTR
dsCycB3#2
dsCycB3#5'UTR
dsCycB3#1
RNAi ctl
RNAi CycB3#1
RNAi CycB3#2
RNAi CycB3 3'UTR
RNAi CycB3 5'UTR
CycB3
Actin
75
50
37
B
% of cells reaching anaphase
RNAi ctl
RNAi CycB3#2
min
NEB
C
% of cells
RNAi ctl
RNAi CycB3#2
DNA bridges
Lagging chromosomes
No anaphase
Multinucleation
D
NEB
RNAi CycB3#2
(SD=38min)
NEB
RNAi ctl
(SD=28min)
Fluorescence intensity
(A.U.)
RNAi ctl
RNAi CycB3#2
min
Anaphase onset

Figure S3

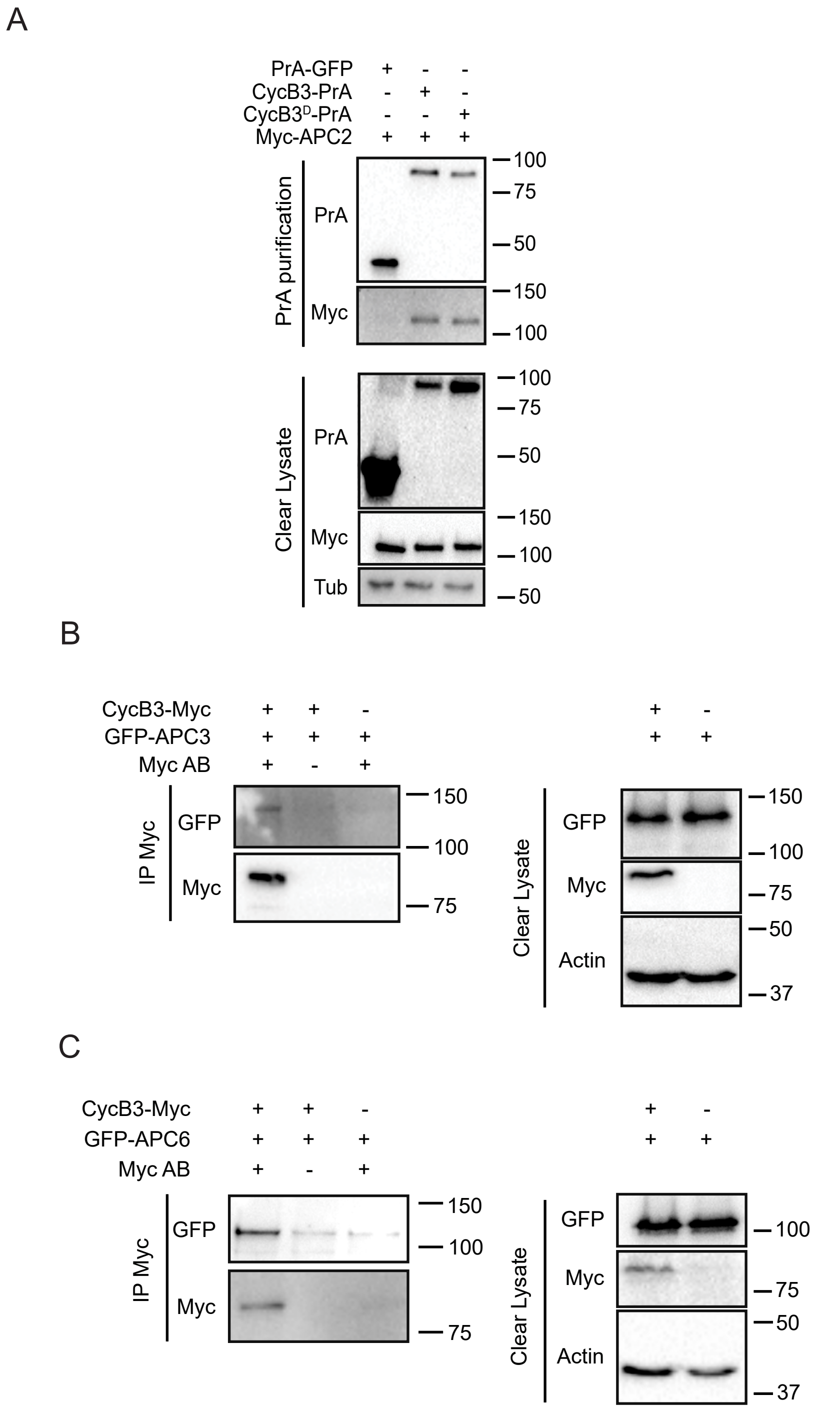
A
PrA-GFP
CycB3-PrA
CycB3D-PrA
Myc-APC2
PrA purification
PrA
Myc
Clear Lysate
PrA
Myc
Tub
100
75
50
150
100
100
75
50
150
100
50
B
CycB3-Myc
GFP-APC3
Myc AB
IP Myc
GFP
Myc
150
100
75
Clear Lysate
GFP
Myc
Actin
150
100
75
50
37
C
CycB3-Myc
GFP-APC6
Myc AB
IP Myc
GFP
Myc
150
100
75
Clear Lysate
GFP
Myc
Actin
100
75
50
37
